# Universal Relation for Life-span Energy Consumption in Living Organisms: Insights for the origin of aging

**DOI:** 10.1101/2020.04.07.030528

**Authors:** Andrés Escala

## Abstract

Metabolic energy consumption has long been thought to play a major role in the aging process (*1*). Across species, a gram of tissue on average expends about the same amount of energy during life-span (*2*). Energy restriction has also been shown that increases maximum life-span (*3*) and retards age-associated changes (*4*). However, there are significant exceptions to a universal energy consumption during life-span, mainly coming from the inter-class comparison (*5, 6*). Here we present a unique relation for life-span energy consumption, valid for ∼300 species representing all classes of living organisms, from unicellular ones to the largest mammals. The relation has an average scatter of only 0.3 dex, with 95% of the organisms having departures less than a factor of *π* from the relation, despite the ∼20 orders of magnitude difference in body mass, reducing any possible inter-class variation in the relation to only a geometrical factor. This result can be interpreted as supporting evidence for the existence of an approximately constant total number N_r_ ∼ 10^8^ of respiration cycles per lifetime for all organisms, effectively predetermining the extension of life by the basic energetics of respiration.

---

Rubner in 1908 (*7*) compared the energy metabolism and lifespans of five domestic animals (guinea pig, cat, dog, cow and horse) and man, finding that the life-span (total) energy expenditure per gram for the five species is approximately constant, suggesting the total metabolic energy consumption per lifespan is fixed, which later has become known as the ‘rate of living’ theory (*1*). Decades later, a mechanism was found in which the idea behind a fixed energy consumption per lifespan might work in the ‘free-radical damage’ hypothesis of aging (*8, 9*), in which macromolecular components of the cell are under perpetual attack from toxic by-products of metabolism, such as free radicals and oxidants. Moreover, energy restriction has been experimentally shown to increase maximum life-span and retard age-associated changes in animals, such as insects, rats, fish, spiders, water fleas and mice (*3, 4*).

Rubner’s relation was confirmed for around hundred mammals (*10*) and extended it to birds (*11*), ectotherms (*12*) and even unicellular organisms such as protozoa and bacteria (*6*), totalizing almost three hundred different species in a range of 20 orders of magnitude in body mass. Although the total metabolic energy exhausted per lifespan per body mass of given organism appears to be relatively constant parameter, at about the same number determined by Rubner of a million Joule per gram of body weight (*7*), variations over an order of magnitude are found among different animal classes, a result previously found by other authors (i.e. *2*) and considered the most persuasive evidence against the ‘rate of living’ theory (*5*).

Recently, the metabolic rate relation was corrected in order to fulfill dimensional homogeneity (*13*), a minimal requirement of any meaningful law of nature (*14*), proposing a metabolic rate (B) formula: 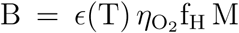, where M is the body mass, f_H_ is a characteristic (heart) frequency, 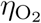 is an specific O_2_ absorption factor and 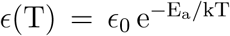 is a temperature correction inspired in the Arrhenius formula, in which E_a_ is an activation energy and k is the Boltzmann universal constant. Compared to Kleiber’s original formulation (*15*), B = B_0_(M*/*M_0_)^0.75^, this new metabolic rate relation has the heart frequency f_H_ as independent controlling variable (a marker of metabolic rate) and the advantage of being an unique metabolic rate equation for different classes of animals and different exercising conditions, valid for both basal and maximal metabolic rates, in agreement with the data in the literature (*13*).

This new metabolic rate relation can be straightforwardly linked (*13*) to the total energy consumed in a lifespan, by the total number N_b_ of heartbeats in a lifetime (N_b_ = f_H_ t_life_), which is empirically determined to be constant for mammals and equals to 7.3 × 10^8^ heart-beats (*16*). Since the total energy consumed in a lifespan relation is valid even in unicellular organisms without heart (*6*), in order to find an unique relation the concept of heart frequency must be generalized to a respiration frequency (f_resp_). This frequency is observed in animals to be proportional to the heart one such f_H_ = a f_resp_ (*17*), which implies that the empirical relation with lifetime is also valid for a total number N_r_ (= N_b_*/*a) of ‘respiration cycles’: 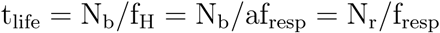

Under the condition of a constant total number N_r_ of respiration cycles in a lifetime t_life_, multiplying the new metabolic rate relation (*13*) by t_life_*/*a = N_r_*/*f_H_ is equivalent to 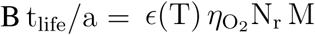. The factor 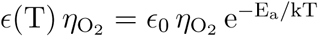 can be rewritten as 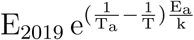, where T_a_ in a normalizing ambient temperature and E_2019_ = 10^−4.313^ mlO_2_g^−1^ ≈ 10^−3^ Jg^−1^ (converting 1 ltr O_2_=20.1 kJ; *17*), comes from the best fitted value for the corrected metabolic relation (*13*). Therefore, for the metabolic rate relation given in (*13*) and under the condition of fixed respiration cycles in a lifetime, the following relation is predicted to be valid:

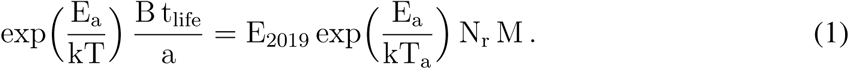

Fig 1 shows the relation predicted by Eq. 1 for 277 living organisms, from unicellular organisms and other ectotherms species, to mammals and birds, listed in Table 1 of (*6*) with their body mass M, total metabolic energy per lifespan B t_life_ and temperature T. The activation energy E_a_ was chosen to the average value of 0.63 eV, independently determined to temperature-normalize the metabolic rates of unicells and poikilotherms to endotherms (*18, 19*). The parameter a was chosen to the empirically determined values of 4.5 for mammals and 9.0 for birds (*17*), 3.75 for ectotherms with heart (estimated from the relative size of their hearts; *17*) and assumed to be unity for ectotherms without heart, such as unicellular organisms. The total number N_r_ = N_b_*/*a = 1.62 × 10^8^ of respiration cycles in a lifetime, was determined from the best fitted values for mammals: N_b_ = 7.3 × 10^8^ heartbeats in a lifetime (*16*) and a=4.5 (*17*). N_r_ will be assumed to be the same for all living organisms from now.

**Fig. 1.**
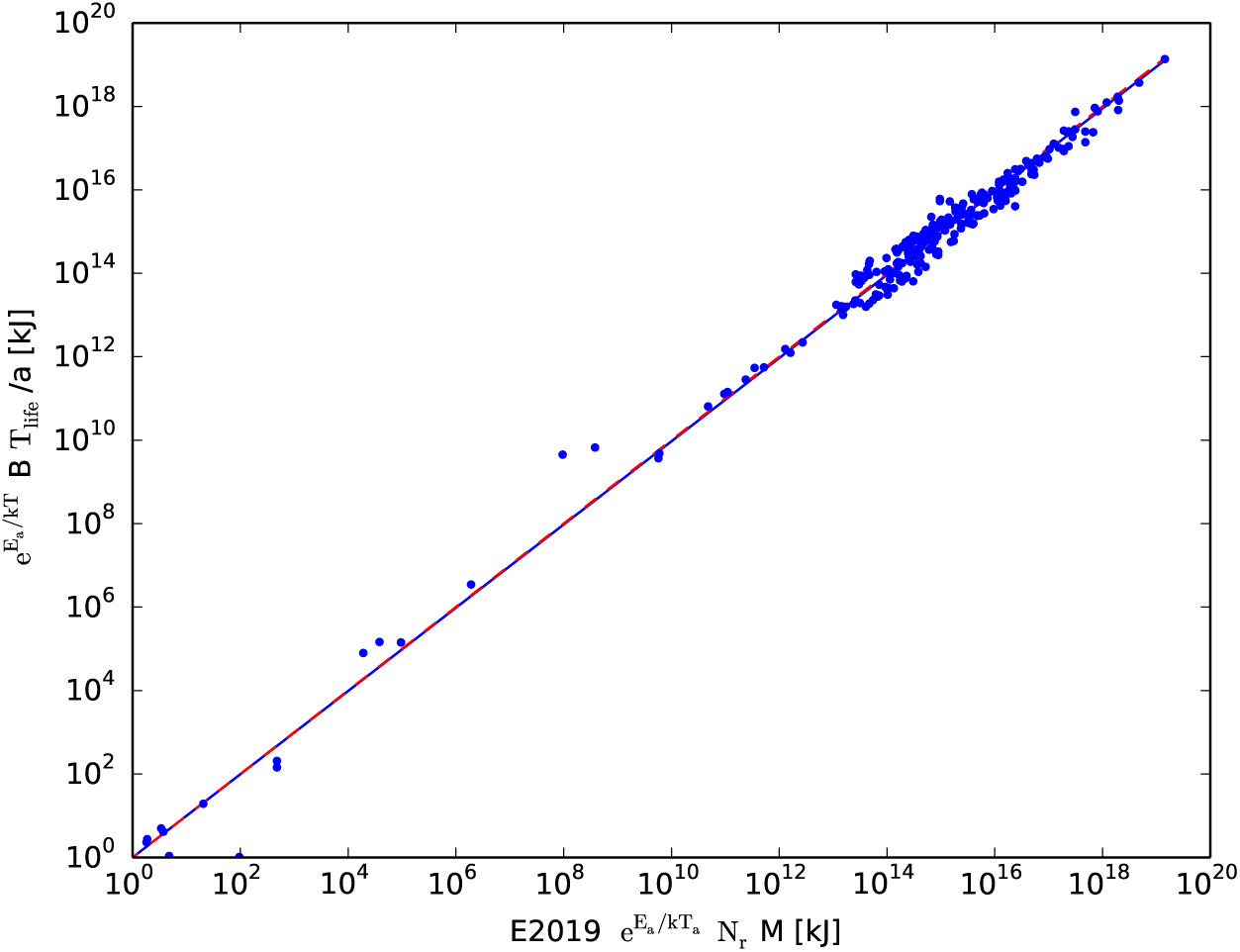
Figure shows the relation predicted by Eq. 1 for 277 living organisms, listed in (*6*). The dashed red curve displays the identity given by Eq 1. The solid blue curve displays the best fitted value of slope 0.997 and normalization of 1.001 for ambient temperature T_a_ = 30°C. The two curves are almost undistinguishable in the dynamical range of 20 orders of magnitude.

The data displayed in Fig 1 support the unique relation predicted by Eq. 1, in a dynamical range of 20 orders of magnitude, for all classes of organisms from *Bacteria* to *Elephas Maximum*. The solid blue curve displays the best fitted value of slope 0.997 and normalization of 1.001, almost undistinguishable from the identity predicted by Eq 1 (dashed red curve). The only free parameter (not predetermined by an independent measurement) in Eq 1 is the ambient temperature T_a_ = 30°C, which was chosen only to match the normalization in the best fitted relation (solid blue) to the identity (dashed red), but is a natural choice for normalization, since ectotherms are typically around 20°C and endotherms close to 40°C in Table 1 of (*6*). Moreover, since the slope close to unity (0.997) is independent of the T_a_ choice, the relation between 5 variables (t_life_, B, a, M & T) given by Eq 1 is confirmed without the choice of any free parameter.

Fig 2 shows the residuals from the relation predicted by Eq 1, as a function of the organism’s body mass. The relation has only an average scatter of 0.339 dex around the predicted value (E_2019_ N_r_ = E_2019_ N_b_/a; dashed line in Fig 2), which impressively small taking into account the 20 orders of magnitude variations in mass and that the values of E_2019_ = 10^−3^ Jg^−1^ (*13*), N_b_ = 7.3 × 10^8^ heartbeats in a lifetime (*16*) and a=4.5 (*17*), comes from three completely independent measurements in mammals. Moreover, around 95% of the points (2 *σ*) has departures from the relation less than a factor of *π* (≈ ±0.5 dex, colored region between the solid curves in Fig 2).

**Fig. 2.**
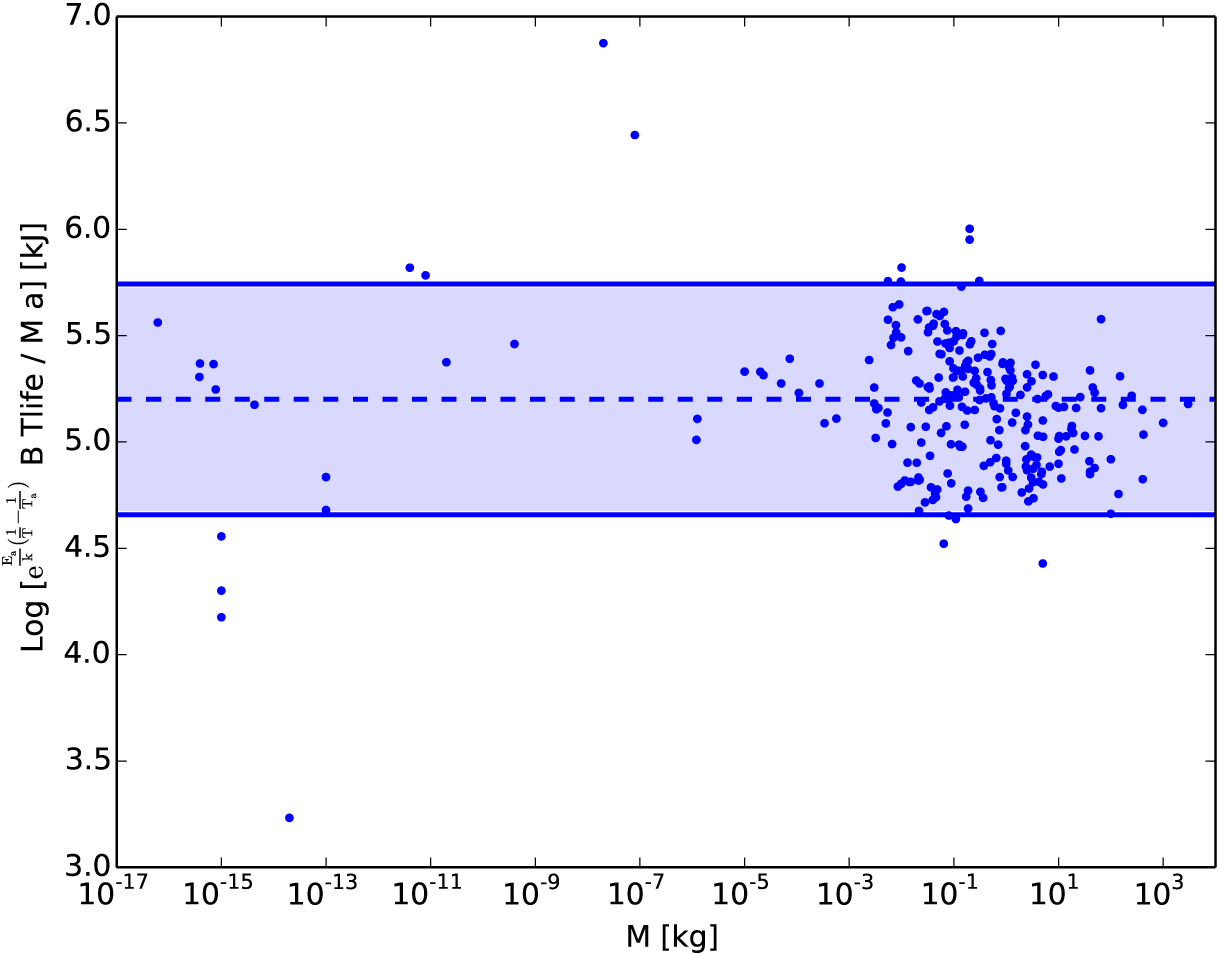
Residuals from the relation predicted by Eq 1 as a function of the organism’s body mass for 277 living organisms, listed in (*6*). The relation has only an average scatter of 0.339 dex around the predicted value (E_2019_ N_r_) denoted by the dashed line. The colored region between the solid curves denotes residuals less than a geometrical factor of *π* from the relation and 95% of the points (2 − *σ*) fulfill such criterion.

The extremely accurate relation displayed in Fig 2, suggests that the only missing parameters are the ones due to geometrical variations among species, of the order of a dimensionless numerical factor order unity (*π*). If that is indeed the case, this is probably the first relation in life sciences including all the relevant controlling parameters, reaching an accuracy comparable to the ones in exact sciences. Moreover, Fig 2 implies no clear trend (larger than an e-fold ≈ 0.5 dex) in inter-class comparison from bacteria to the largest mammal.

Inter-class variations was considered the most persuasive evidence against the ‘rate of living’ theory, for example, the inter-class comparison of birds and bats to mammals (*2, 5*). These exceptions were erased in Fig 2 due to predicted secondary parameters that are not included in the original formulation (mainly the parameter a = f_H_*/*f_resp_ for the particular case of birds and bats). Moreover, Eq 1 implies a dependence of 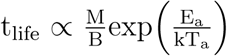 that it is in agreement with the three regimens experimentally known to extend life-span (*20*): lowered ambient temperature T_a_ in poikilotherms, decrease of physical activity in poikilotherms (lower B) and caloric restriction (lower B).

Invariant quantities in physics traditionally reflects fundamental underlying constraints, something recently also applied to life sciences such as Ecology (21, 22). Fig 2 displays the fact that, for a given temperature, the total life-span energy consumption per gram per ‘generalized beat’ 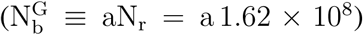 is remarkably constant on around E_2019_, supporting that the overall energetics during lifespan is the same for all living organisms, being predetermined by the basic energetics of respiration. Therefore, Rubner’s original picture it is shown to valid without systematic exceptions, but in this more general form. In addition, we showed here that this invariant comes directly from the existence of another invariant: the approximately constant total number N_r_ ∼ 10^8^ of respiration cycles per lifetime, effectively converting the ‘generalized beat’ into the characteristic clock during lifespan. Thus, the exact physical relation between (oxidative) free radical damage and the origin of aging, is most probably related to the striking existence such of constant total number of respiration cycles N_r_ in the lifetime of all organisms, which predetermines the extension of life.

It have been also suggested that an analogous invariant is originated at the molecular level (23), the number of ATP turnovers in a lifetime of the molecular respiratory complexes per cell, which from an energy conservation model that extends metabolism to intracellular levels is estimated to be ∼ 1.5 × 10^16^ (23). Similar number can be determine taking into account that human cells requires synthase approximately 100 moles of ATP daily, equivalent to 7 × 10^20^ molecules per second. For ∼ 3 × 10^13^ cells in human boby and for a respiration rate of 15 breaths per minute, this gives ∼ 9 × 10^7^ ATP molecules synthesized per cell per breath, which for the invariant total number N_r_ of respiration cycles per lifetime found in this work, arises to the same number of ∼ 1.5 × 10^16^ ATP turnovers per cell in a lifetime.

Finally, the empirical support in favor of Eq 1 allow us to compute how much will vary the energy consumption in the biomass doing aerobic respiration, as increases the earth’s ambient temperature T_a_, relevant in the current context of possible global warming. This is given by the factor 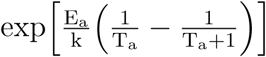 which for an activation energy E_a_ = 0.63eV and ambient temperature T_a_ = 30°C, implies an increase 8.3% in energy consumption per 1 degree increase on the average Earth temperature T_a_. This result can be straightforwardly applied in endotherms since their body temperatures adapt to the environmental one, however, is less clear its implications for the case of ectotherm organisms.

## Acknowledgments

I acknowledge partial support from the Center of Excellence in Astrophysics and Associated Technologies (AFB-170002) and FONDECYT Regular Grant 1181663.

